# MitoSearch: Construction of fish species composition database with environmental DNA using public databases

**DOI:** 10.1101/2022.03.09.483700

**Authors:** Takumi Ito, Hideaki Mizobata, Kazutoshi Yoshitake, Shigeharu Kinoshita, Shuichi Asakawa

**Affiliations:** Laboratory of Aquatic Molecular Biology and Biotechnology, Aquatic Bioscience, Graduate school of Agricultural and Life Sciences, The University of Tokyo

## Abstract

Existing fish species distribution databases have the following problems: the possibility of species identification errors due to plasticity of morphological characteristics, the impossibility of quantitative composition, and the difficulty of geographically and time-series integrated interpretation. To solve the above problems, we constructed MitoSearch database. It is a fish species distribution database using 2,160 samples of variable regions of 12S rRNA genes amplified by MiFish primers registered at NCBI/EBI/DDBJ, and visualizes the fish species composition data on a map. By integrating a large amount of data registered in public databases, MitoSearch was aimed to provide the comprehensive understanding of spatial and temporal distribution of organisms on a global scale.

## 1. Introduction

### 1-1. species identification by DNA barcoding

In 2003, the DNA barcoding method was established for species identification using the mitochondrial gene cytochrome c oxidase (CO1). Prior to this, species identification was based solely on morphological characteristics. However, species identification based on morphological characteristics could lead to incorrect species identification due to morphological plasticity caused by genetic variation, sex differences, and life stages. Therefore, the DNA barcoding method was established to identify species by amplifying barcodes consisting of highly conserved primer regions and species-specific gene regions and decoding their sequences. It was also reported that the CO1 gene, which has a fast rate of molecular evolution and a highly conserved region among species, has specific sequences in 95% of animal species, and that DNA barcoding using the CO1 gene as a barcode is very effective [1].

### 1-2. eDNA barcoding for abundance estimation

Genetic material left behind by organisms in non-living components of the environment (soil, sediment, water, etc.) is called environmental DNA (eDNA) [2]. In the past, targeting fragments of less than 100 bp of the mitochondrial gene cytochrome b (cytb), the eDNA barcoding technique has been shown to be applicable in seawater samples, allowing eDNA barcoding to detect the same fish species found in a catch, and detection of saltwater fish by eDNA barcoding has proven to be effective [3]. Furthermore, it has been reported that there is a positive correlation between the abundance of fish species in environmental water and the amount of eDNA obtained from environmental water, and that eDNA barcoding can be used not only to detect fish species but also to estimate the abundance of fish species [4][5]. Abundance estimation with eDNA barcoding has three advantages over abundance estimation with conventional fishing: it has less impact on the environment and ecosystems because it does not involve fishing, it can detect rare species, and it does not introduce errors in species identification due to morphological plasticity [6]. These advantages, coupled with the advent of next-generation sequencers, have led to the current popularity of eDNA metabarcoding for abundance estimation.

### 1-3. understanding fish species composition with MiFish primers

MiFish, a primer for metabarcoding fish environmental DNA (eDNA), was developed in 2015 [7]. This primer targets the variable region of the 12S rRNA gene (163-185 bp). This MiFish primer has been reported to detect more fish species than primers used in other eDNA barcoding [8]. Sequencing data of 12S rRNA genes amplified by MiFish primers can also provide information on the composition of fish species in environmental water through the MiFish pipeline [9].

### 1-4. Existing fish stock databases and their challenges

In 1996, fish distribution and abundance were surveyed in medium to large 38 rivers in New Zealand, leading to the development of a database of fish distribution in New Zealand [10]. In 2007, RSD, a database containing morphological, ecological, and distributional information on fish species in Taiwan was constructed [11]. In 2013, FiMSEA, a database integrating taxonomic, distributional, and photographic records of freshwater fish specimens in mainland Southeast Asia collected by several research institutions and its visualization platform was constructed [12]. However, the current fish species distribution database and its visualization platform have the following three problems: 1. the possibility of incorrect species identification and missing rare species cannot be eliminated because the fish species distribution is based on identification with collection and morphology, 2. only the distribution of existing fish species can be known, and 3. However, the current fish species distribution database and its visualization platform have the following three problems: the possibility of incorrect species identification and missing rare species cannot be eliminated because the fish species distribution is based on identification by collection and morphology, only the distribution of existing fish species can be known, and the information can only be obtained for each sample data unit, making geographically and time-series integrated interpretation difficult.

### 1-5. Objectives and outline of this study

In order to solve the above problem and to enable geographically and time-series integrated interpretation of fish species composition in various water bodies on an unprecedented scale, we constructed a fish species composition database by extracting variable regions of 12S rRNA genes amplified by MiFish primers from next-generation sequencing data registered at NCBI, and developed a platform of MitoSearch to map fish species composition data, integrate data across samples, and understand time series.

## 2. Method

### 2-1. Acquisition of 12S rRNA gene sequencing data amplified by MiFish primers

SRA numbers were searched and obtained from the NCBI SRA database (https://www.ncbi.nlm.nih.gov/sra) with the keyword of “mifish”. The date of collection and the latitude and longitude of the sampling point were described in SRA sample information. The sequencing data corresponding to each SRA number were then downloaded in gzipped FASTQ format using sratoolkit version 2.11.1.

### 2-2. Data preprocessing

The paired-end sequencing reads of the downloaded sequencing data were jointed by using FLASh. The -M parameter was specified as 300 to set the maximum overlap length to 300 bp. Single-ended sequencing data were used without processing by FLASh. The FASTQ files were then converted to FASTA files.

### 2-3. Homology search to genome database

A homology search was performed using blastn against the MitoFish database using the above FASTA files as queries. The -max_target_seqs 1 option was specified to extract only the most homologous species for each read. We then removed hits with less than 90% homology, removed hits that were only aligned in areas less than 90% of the read length, and removed reads that were less than 100 bp, as the region amplified by the MiFish primers is approximately 200 bases. 17 samples were randomly selected, then MitoFish and nt homology searches were performed and compared in the databases. The nt database was downloaded on May 21, 2021. The percentage of reads hit in the MitoFish database that were non-fish in the nt database was calculated before and after filtering.

### 2-4. Construction of fish species composition database

After filtering was completed, the number of hit reads was calculated for each fish species in the blast search results. Each fish species was assigned a Japanese name based on the list of Japanese fish species (https://www.museum.kagoshima-u.ac.jp/staff/motomura/20210718_JAFList.xlsx). Samples with less than 100 reads in the total composition were removed as they were considered to be greatly affected by noise, and fish species that accounted for less than 1% of the total composition were removed to reduce noise. The composition of each fish species was then adjusted so that the composition was 100 in total for all fish species, and the sampling date and time were assigned. Finally, data from the same species with different Accession IDs were merged.

### 2-5. Construction of MitoSearch, a platform for utilizing the fish species composition database

OpenStreetMap was used for the Map. D3.js, a web graphing library, was used to draw the graph. Leaflet library was used to draw the graph on the map. Marker Cluster Group, an extension library of Leaflet, is used to integrate geographically adjacent samples.

### 2-6. Analysis using MitoSearch

Ridge graphs showing the temporal abundance trends of each fish species in Tokyo Bay in MitoSearch were used for analysis.

## 3. Results

### 3-1. Acquisition of next generation sequencing data with samples using MiFish primers

We were able to obtain 4,355 samples registered as amplified by MiFish primers from the next-generation sequencing data registered at NCBI/EBI/DDBJ.

### 3-2. Homology search to the genome database

The MitoFish database was selected because it has a sufficient number of fish 12S rRNA genes registered in MitoFish (36,967 species), and the data size is smaller than the NCBI nt data, allowing for faster homology searches. However, when conducting this homology search, since only fish are registered in the MitoFish database, there is a possibility that there are reads that were hit as fish in the MitoFish database but were not originally derived from fish. We checked the nt database for non-fish reads and found that there were many reads with a read length of about 50 bp. These reads may have originated from the adaptor sequence of the illumina. Therefore, to confirm that reads that hit as fish in the MitoFish database but were not originally derived from fish were removed by filtering more than 100 reads, 90% match rate, and 90% coverage, 17 samples were randomly selected, then MitoFish and nt homology searches were performed and compared in the databases. The percentage of reads hit in the MitoFish database that were non-fish in the nt database was calculated before and after filtering. The percentage of reads that hit the nt database among the reads that hit the MitoFish database before and after filtering changed from 7.3% to 0% (Table 1). This indicates that our filtering eliminated reads that did not originate from fish effectively.

**Table 1.** Number of reads hit by MitoFish before and after filtering that are non-fish in the nt database.

|  | Before<br>Filtering | After<br>Filtering |
| --- | --- | --- |
| Average number of reads that hit the MitoFish database [reads] | 28304 | 22327 |
| Average number of non-fish hits in nt database among reads hit in MitoFish database [reads] | 2069 | 0 |
| Percentage of reads that hit the nt database among the reads that hit the MitoFish database [%] | 7.3 | 0 |

### 3-3. Construction of fish species composition database

From 4,355 eDNA sequencing data, we were able to construct a fish species composition database consisting of 2,559 composition data (Table 2). This means that 1,796 eDNA sequencing data were lost in the process of database construction. About 800 data were removed due to the fact that the sample registered one read in the Sanger sequence. The compositional database allowed an average of 6.64 fish species to be detected per sample, for a total of 827 fish species in all samples. Of the 2,559 compositional data, 2,560 samples contained latitude and longitude information (Table 2). The average number of reads of the downloaded sequencing data was 90,934 reads, and the average number of reads lost after FLASh joint was 22,759 reads. The average number of reads that did not hit the MitoFish database was 23,168 reads, and the average number of reads lost after filtering was 17,630 reads (Table 3).

**Table 2.**
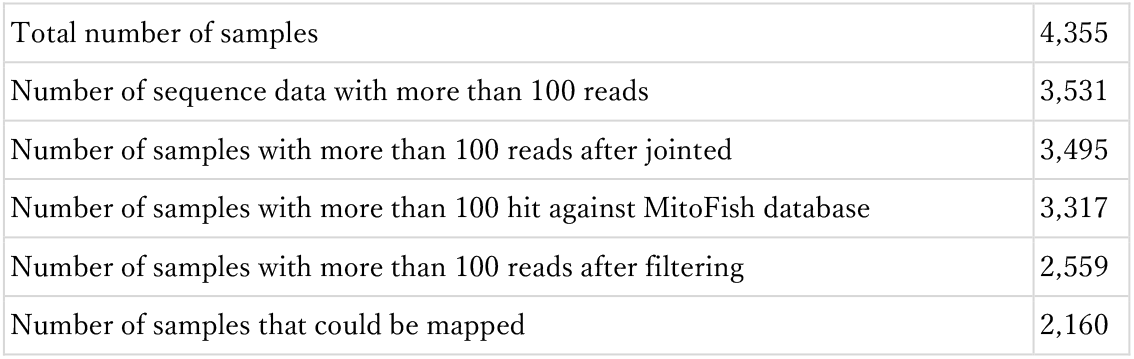
Number of samples in the process of database construction

|  |  |
| --- | --- |
| Total number of samples | 4,355 |
| Number of sequence data with more than 100 reads | 3,531 |
| Number of samples with more than 100 reads after jointed | 3,495 |
| Number of samples with more than 100 hit against MitoFish database | 3,317 |
| Number of samples with more than 100 reads after filtering | 2,559 |
| Number of samples that could be mapped | 2,160 |

**Table 3.** Average number of reads in the process of database construction

|  |  |
| --- | --- |
| Average number of reads for raw sequencing data | 90,934 |
| Average number of reads after FLASH joint | 68,175 |
| Average number of hits against MitoFish database | 45,007 |
| Average number of reads after filtering | 27,377 |

### 3-4. Construction of MitoSearch, a platform for utilizing fish species composition database

We implemented the following four main functions of MitoSearch, an online fish composition database visualization platform.

#### 3-4-1. Mapping the composition of each sample on a map

For each fish species composition data including latitude and longitude data in the database, D3.js is used to draw a pie chart displaying the composition. These pie charts are plotted as markers at the same latitude and longitude as the sampling points on the map (Fig 1). These pie charts also have tooltips that can be hovered over to display the sample’s SRA number, collection date, and species composition (Fig. 2). The pie charts also have a pop-up function that allows the user to click to display the sample’s SRA number, collection date, and species composition, as well as a link embedded in the SRA number to access a NCBI web page with sample information.

**Fig 1.**
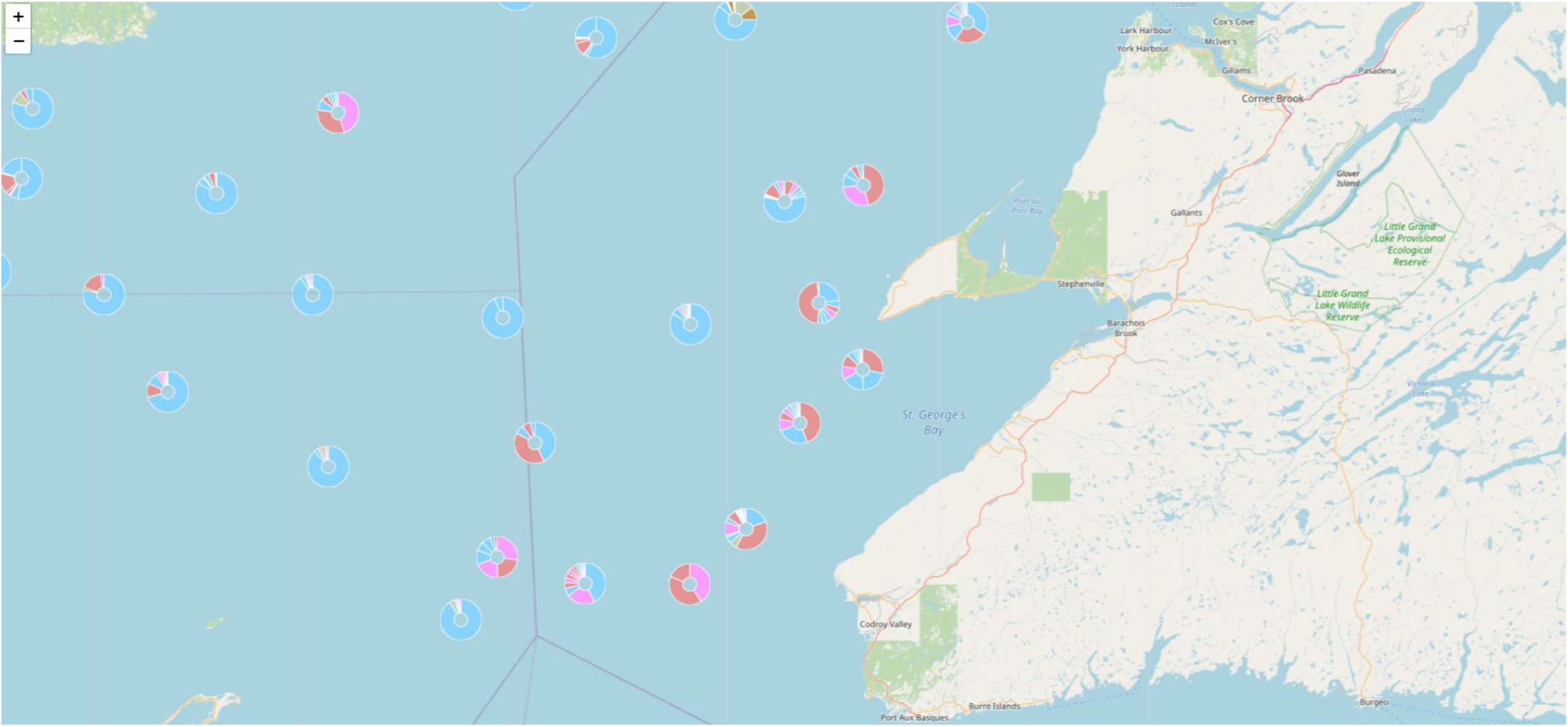
Pie chart showing fish species composition mapped by MitoSearch. A pie chart showing fish species composition is mapped at the same latitude and longitude as the point where the sample was taken. The coloring is done according to the taxonomic group of the fish species.

**Fig 2.**
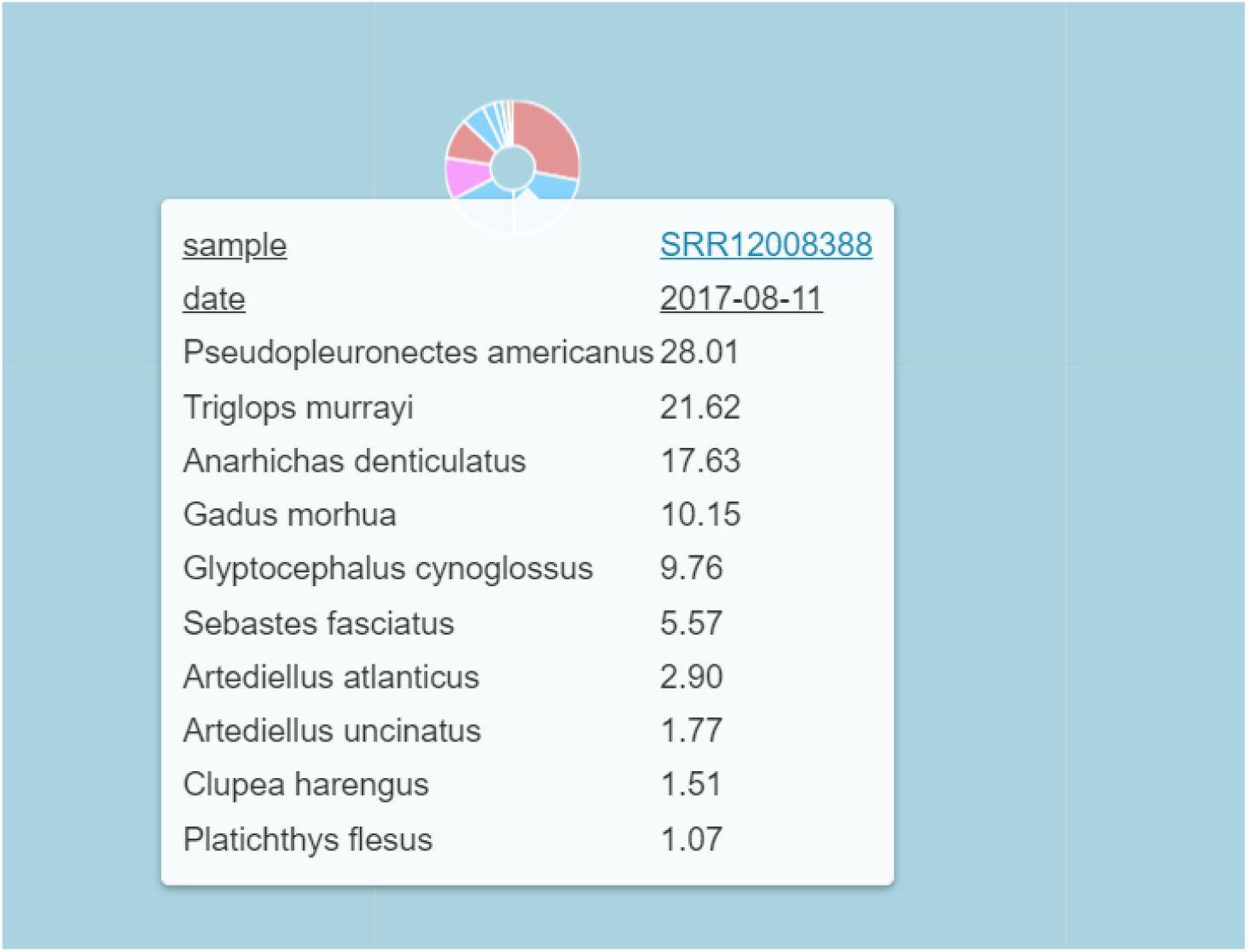
Tooltip feature to display the specific fish composition of a sample in MitoSearch. By hovering the mouse over the sample, the SRA number, collection date, and species composition of the sample can be displayed. In addition, the SRA number, collection date, and species composition of the sample can be displayed by clicking, and a link embedded in the SRA number can be used to access NCBI’s Web page with sample information.

#### 3-4-2. Integration of geographically close samples

Geographically adjacent samples are integrated as the same cluster according to the map display area, and the fish species composition of the entire cluster is plotted as a pie chart (Fig. 3). The composition of the entire cluster is calculated for each fish species by accumulating the percentage of the composition of the relevant fish species in the child elements, and finally averaging so that the composition is 100 for all fish species. The radius of this pie chart increases with the number of subelements in the cluster. By clicking on the pie chart, the cluster can be split and the child clusters can be displayed or merged again. The pie chart also has a tooltip function that allows the user to mouse over the pie chart to display the number of samples in a cluster and the averaged composition of the top 20 fish species in the cluster.

**Fig 3.**
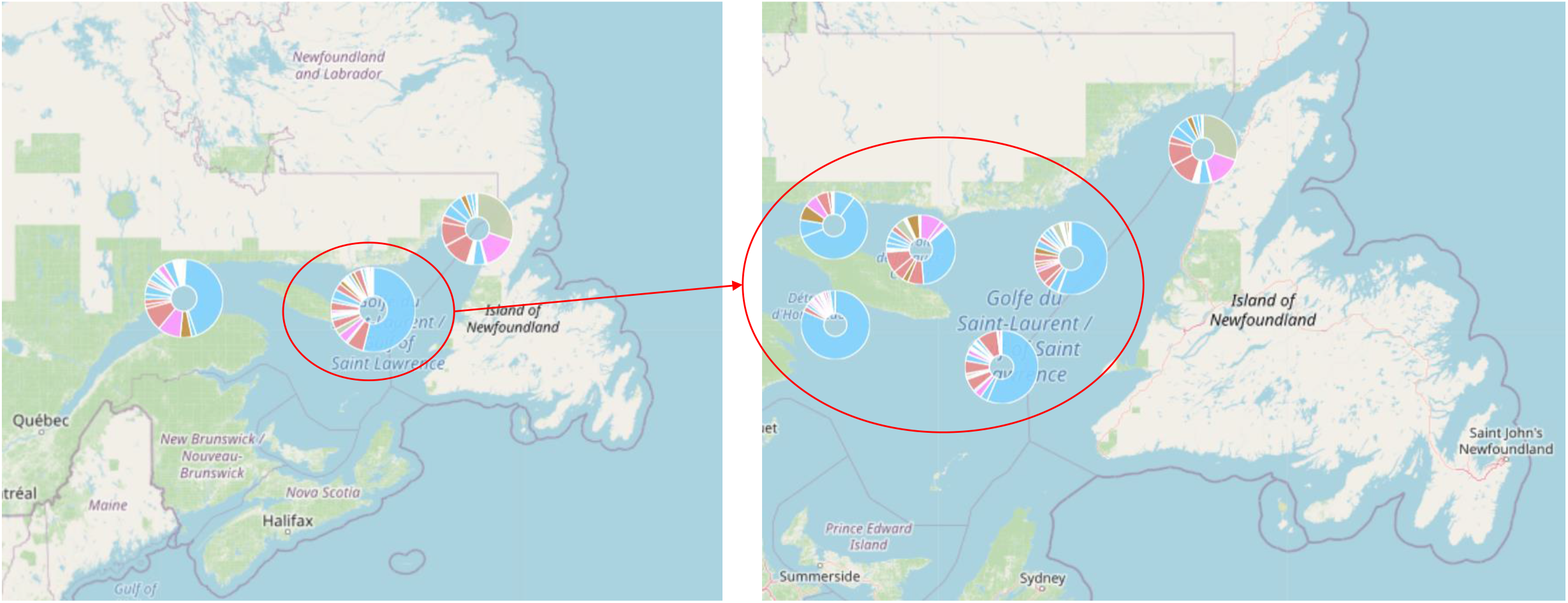
Pie chart showing fish species composition merged with geographically close samples in MitoSearch and tooltip feature showing fish composition. The composition of the entire cluster is calculated by summing the percentage of the composition of the relevant fish species in the child elements for each fish species, and finally averaging the compositions for all fish species to reach 100. This pie chart also has a larger radius as the number of sub-elements in the cluster increases. By clicking on the pie chart, the cluster can be split and the child clusters can be displayed or merged again. The pie chart also has a tooltip function that allows the user to mouse over the pie chart to display the number of samples in a cluster and the averaged composition of the top 20 fish species in the cluster.

#### 3-4-3. Visualization of temporal abundance trends

For each fish species present in the composition within the map display area, a ridge graph displays how the abundance within the map display area changes (Fig. 4). The height of the ridge graph is adjusted to be constant for all fish species. The graph becomes darker as the percentage of the composition at the point of maximum abundance increases.

**Fig. 4.**
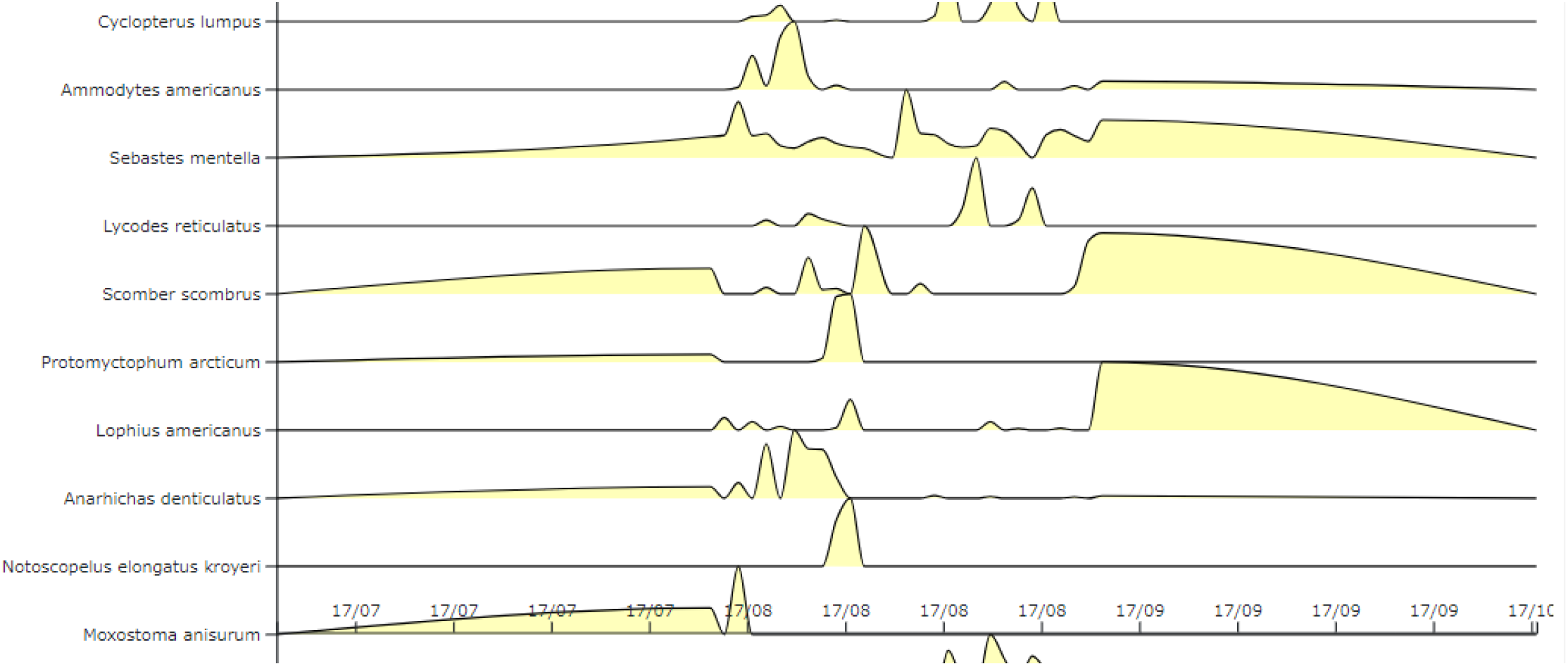
Ridge graph showing time trends of abundance within the displayed area of the map in MitoSearch. The height of the ridge graph is adjusted to be constant for all fish species. The graph becomes darker as the percentage at the point of maximum abundance increases.

#### 3-4-4. Filtering samples by species name

By entering the name of a fish species in the text-entry select box, the pie and ridge graphs can be narrowed down to samples that contain only the species entered (Fig. 5). Multiple fish species can also be selected. All data except for the fish species used for filtering are accumulated as “others”.

**Fig. 5.**
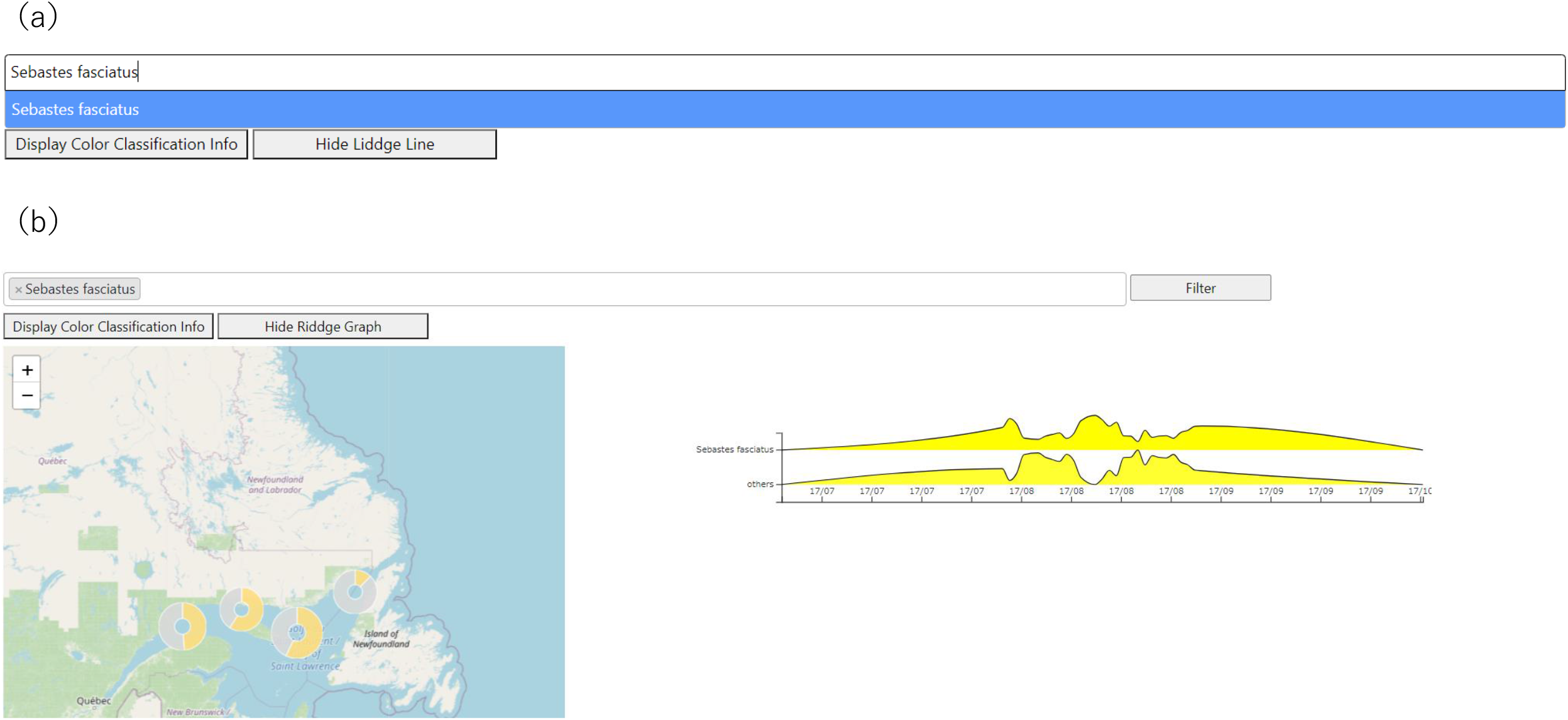
Filtering function in MitoSearch to narrow down the display of composition to specific fish species. By entering the name of a fish species in the text-entry select box, it is possible to display a pie chart or ridge graph by narrowing down the sample to those containing only the species entered. (a) Text input box for entering the species of fish you wish to filter. (b) A pie chart showing the composition of fish species after filtering.

### 3-5. Analysis using MitoSearch

MitoSearch data on time trends of fish abundance in Tokyo Bay revealed that sea bass, *Lateolabrax japonicus* accounted for a high proportion of the fish composition during the winter season (Fig. 6). This is consistent with previous literature describing the migration of sea bass from rivers to offshore areas for spawning in winter [13][14].

**Fig 6.**
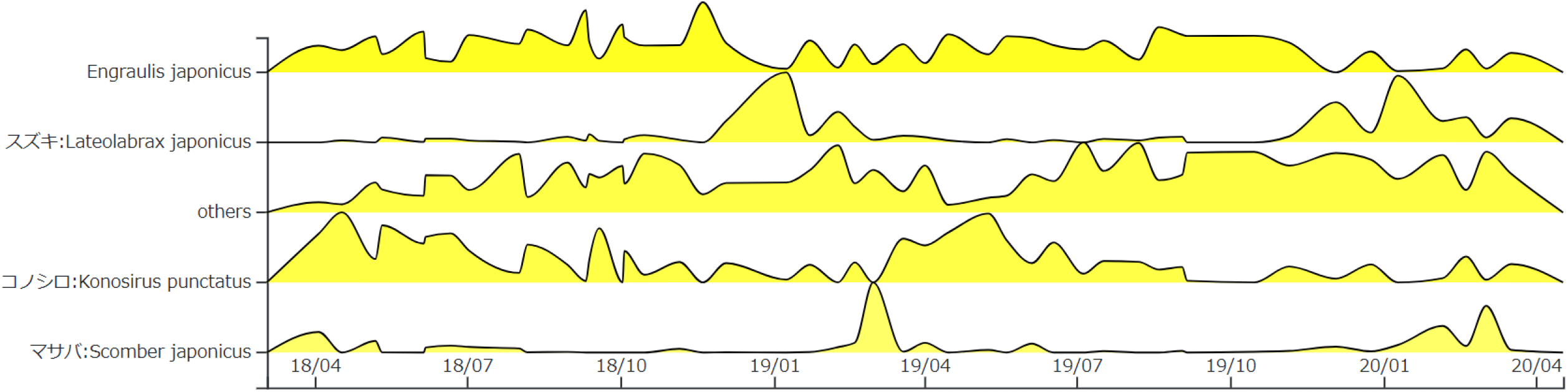
Data obtained by MitoSearch on fish abundance trends in Tokyo Bay. Abundance trend of Japanese anchovy, sea bass, whitebait, and chub mackerel is shown.

The data also revealed that *Konosirus punctatus* made up a high proportion of the species composition in April and May (Fig. 6). This finding is in agreement with previous literature indicating that *K. puncttatus* spawn in April and May [15].

Furthermore, Japanese anchovy, *Engraulis japonicus* was observed to be highly represented in the species composition of Tokyo Bay throughout the year (Fig. 6). This is consistent with previous literature describing the high year-round catch of Japanese anchovy in plankton-rich Tokyo Bay [16]. This agreement with data from the past literature indicates the usefulness of this tool.

The same data showed that chub mackerel, *Scomber japonicus* accounted for a high percentage of the species composition from March to April (Fig. 6). In the past literature, it has been reported that the spawning season of chub mackerel depends on sea water temperature, and that the spawning season of chub mackerel in the East China Sea extends from February to June [17]. Existing literature has not reported seasonal trends in abundance or spawning seasons of chub mackerel in Tokyo Bay. Therefore, this study is the first to suggest that the abundance of chub mackerel in Tokyo Bay is high from March to April. However, it is not clear which growth stage of the mackerel stock is abundant on this platform, and further research is needed to determine why the mackerel are abundant in Tokyo Bay from March to April. However, it is known that the amount of eDNA in the environment varies depending on various factors such as species morphology, water temperature, and currents [18][19][20]. Careful consideration needs to be given regarding the method used to calculate fish species composition.

## 4. Discussion

In this study, fish species composition was calculated by counting the best hit reads for each fish species. Recently, mtDNA 16S RNA genes have been targeted to detect bivalves and gastropods in lakes [21]. In the future, it will be possible to integrate this information into the database to build a database of aquatic biota composition that includes not only fish but also invertebrates.

In this study, there were 399 composition data that could not be mapped because the latitude and longitude information of the samples was not provided. We hope that by promoting the usefulness of this database, sequencing data will be registered in a correct format. Continuous eDNA sampling is currently limited to a few regions, but if eDNA sampling were done on many countries continuously, this database could provide more valuable information.

## 5. Conclusion

We constructed MitoSearch database. It is a fish species distribution database using 2,160 samples of variable regions of 12S rRNA genes amplified by MiFish primers registered at NCBI/EBI/DDBJ, and visualizes the fish species composition data on a map. By integrating a large amount of data registered in public databases, MitoSearch was aimed to provide the comprehensive understanding of spatial and temporal distribution of organisms on a global scale. The data obtained from this platform on abundance trends of sea bass, whitebait, and Japanese anchovy in Tokyo Bay were consistent with previous literature that investigated abundance by sampling and other methods, demonstrating the usefulness of this platform. The tool also revealed that the proportion of chub mackerel in Tokyo Bay was high from March to April. In the future, it will be possible to construct a database of aquatic organism composition including invertebrates by integrating environmental DNA information using universal primers other than fish. If environmental DNA studies are conducted continuously over a wider area and sequence data are registered in public databases, this database will become the organism distribution database that will provide more valuable information.

## 6. Software and code availability

MitoSearch can be accessed at the following link. (https://meta.fs.a.u-tokyo.ac.jp/mitosearch/)

